# Sealable capped nanovials for high-throughput screening of cell growth and function

**DOI:** 10.1101/2025.06.29.662236

**Authors:** Michael Mellody, Yuta Nakagawa, Alyssa Arnheim, Lily Shang, Catherine Soo, Natalie Tsubamoto, Sarah Taylor, Sumedha Shastry, Wesley Luk, Ian Morales, Richard James, Dino Di Carlo

## Abstract

Millions of modular nanoliter-scale compartments that isolate functionally rich single-cell and cell-to-cell communication data can scale biological discovery for the age of AI. Here, we introduce capped nanovials—suspendable, sealable microscale compartments formed by the docking of hydrogel capping particles into bowl-shaped nanovials—as a versatile system for culturing, analyzing, and sorting single cells and small colonies. This two-particle architecture enables localized confinement of cells and secreted products while maintaining compatibility with standard laboratory workflows such as wash and reagent exchange steps, fluorescence microscopy, and flow cytometry. Crucially, these compartments are formed via simple pipetting and centrifugation steps, making the platform highly democratized. We demonstrate the ability of capped nanovials to compartmentalize single mammalian, bacterial, and yeast cells and support growth into colonies, enabling selection based on proliferation and bioproduction. We further show that capped nanovials enhance single-cell secretion assays by reducing molecular crosstalk and increasing signal-to-noise ratios. Importantly, we demonstrate functional co-culture assays by permitting stable confinement of cell pairs, enabling detection and enrichment of antibody-secreting cells based on the ability of their secreted antibodies to activate co-encapsulated reporter T cells, achieving a signal-to-noise ratio of >30 and up to 100% selection purity. By combining the simplicity of standard lab handling with the resolution and throughput of traditional microfluidic compartmentalization approaches, capped nanovials provide a new class of scalable, accessible test tubes for modern single-cell biology.

## Introduction

Throughout the history of biological research, experimentation has relied on a simple yet powerful concept: placing cells or reagents into a defined compartment or vessel. From the petri dish and test tube to the well plate and growth flask, these vessels have enabled observation, growth, and manipulation of biological systems. As biological inquiry has pushed toward the resolution of individual cells and molecules, the vessels for experimentation are evolving—miniaturizing toward a "quantum" limit where each unit contains a single cell or a microcolony, yet remains accessible, manipulatable, and scalable. To meet the demands of modern biology and generation of huge datasets to train AI models, these next-generation test tubes should support millions of individual experiments simultaneously,^1^ allow analysis by standard laboratory instruments, and enable characterization of the long-term dynamics of living cells.^2^ Microfabricated well arrays^3–7^, valve-based microfluidics,^8^ and droplet microfluidics^9,10^ have provided early attempts at this transition, yet often require specialized infrastructure, do not enable long-term culture, or trade throughput for flexibility.^11^ The first generation of lab-on-a-particle technologies,^12^ such as hydrogel nanovials^13,14^ and core-shell particles,^15,16^ have begun to address scalability, long-term culture without nutrient depletion, and accessibility, but remain limited by open architectures or by fabrication methods that demand expertise in microfluidics.^17^ Here, we introduce capped nanovials as a new class of sealable, suspendable test tubes for single cells, cell pairs, and cell-derived colonies. These microscale compartments can be loaded, sealed, incubated for cell culture while enabling nutrient/waste exchange, and analyzed using only standard laboratory equipment, enabling massively parallel assays for growth, secretion, and cell-cell interactions. By combining the simplicity of traditional macroscale vessels with the scale and control required for single-cell science, this platform reimagines the vessels for biological experimentation—opening new avenues for accessible, high-throughput discovery.

## Results

### Generating suspendable compartments through complementary docking of shaped particles

To create a microscale assay platform capable of analyzing and isolating single cells and multi-cell interactions in a high-throughput manner, we developed a two-particle interlocking system composed of larger diameter bowl-shaped hydrogels (nanovials) and smaller diameter spherical hydrogels (capping particles) (Figure 1A). Nanovials are hydrogel microparticles with cavities that can be functionalized with biomolecules and sized to hold cells.^13,14^ Through mixing and centrifugation processes we found that smaller capping particles are able to stably dock within the cavities of the nanovials, capping the nanovials and creating a multitude of nanoliter-scale volume compartments surrounded by the hydrogel materials of the nanovial and capping particle. These “capped nanovials” impart unique functions to the compartment depending on how each component (the nanovial and capping particle) is chemically modified to attach proteins, antigens, antibodies and other molecules (Figure 1A). A standard workflow enables encapsulation of cells within capped nanovials, which can be observed through standard microscopy (Figure 1B) or flow cytometry, as described later. Thousands to millions of capped nanovials can be formed in parallel in solution creating sealed compartments for clonal cell culture, on-particle assays for growth or secretion, and cell-cell interaction studies (Figure 1C).

**Figure 1.**
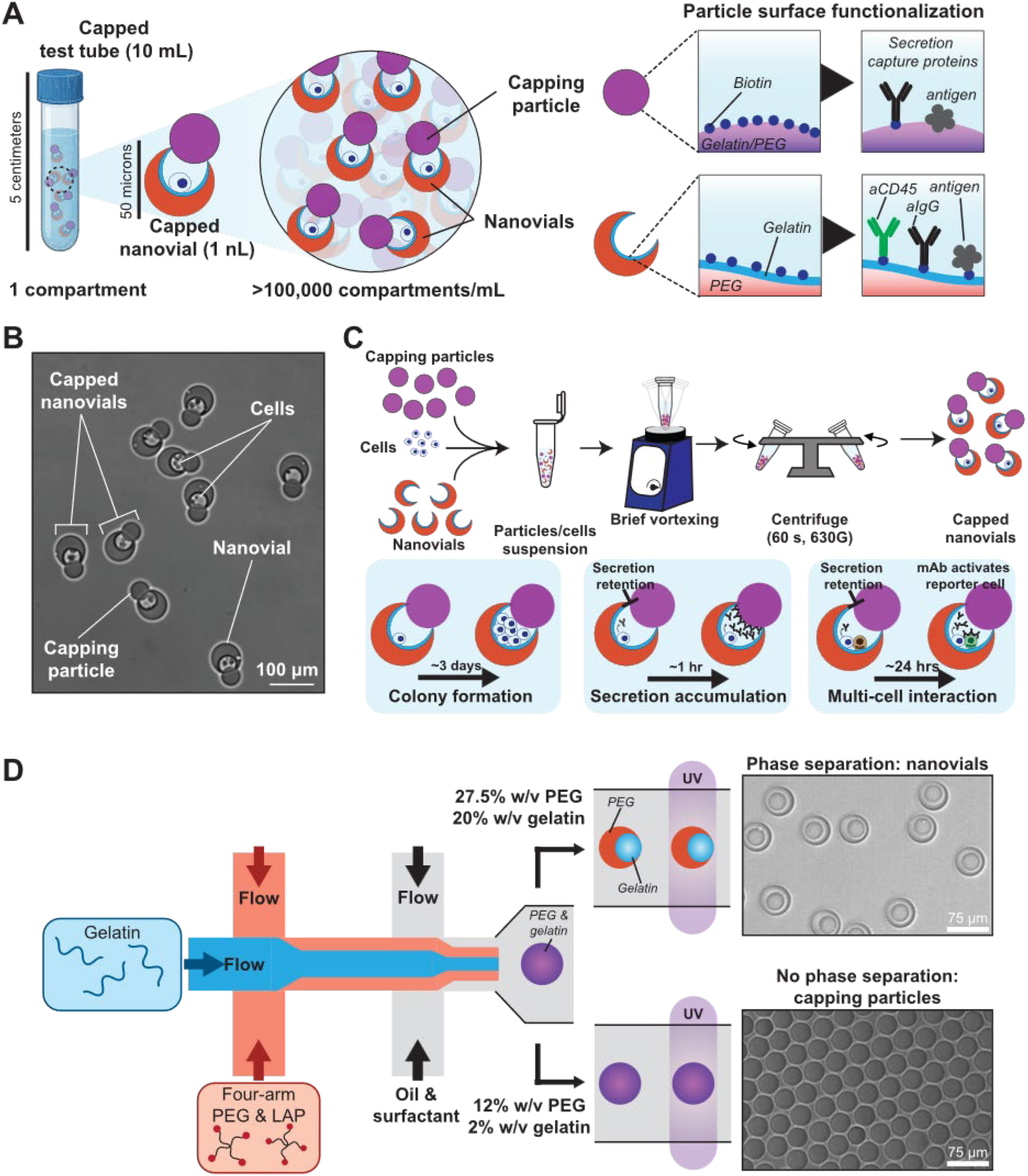
Capped nanovial workflow for single-cell assays. **A**) Capped nanovials are likened to sealable, microscopic test tubes composed of two multi-functional hydrogel particles that can be decorated with proteins and other chemical moieties. **B)** Representative brightfield image of capped nanovials loaded with mammalian cells. **C)** Cells are loaded into nanovials, followed by centrifugation with spherical capping particles in order to create sealed, solid-phase reaction vessels for downstream applications such as growth studies, secretion accumulation and detection, and multi-cell interaction assays. **D)** Nanovials and capping partic les are fabricated using an aqueous two-phase system composed of a gelatin stream and a polyethylene glycol (PEG) and lithium phenyl-2,4,6-trimethylbenzoylphosphinate (LAP) stream. Reducing PEG and gelatin concentrations limits phase separation, resulting in production of capping particles rather than nanovials.

Both nanovials and capping particles of varying sizes are fabricated using a microfluidic droplet generator of water-in-oil droplets comprising a polyethylene glycol (PEG)-gelatin aqueous two-phase system (Figure 1D).^13^ Phase separation of PEG and gelatin is concentration dependent, following a binodal curve.^13^ At high concentrations above the binodal curve, PEG and gelatin phase separate to create bowl-shaped nanovials after ultraviolet (UV) crosslinking, whereas at low concentrations below the binodal curve phase separation does not occur, resulting in spherical capping particles. We adjusted the microfluidic device dimensions to manufacture nanovials and capping particles of varying sizes. Nanovials of two sizes were manufactured for use in this work, ∼40 µm and ∼70 µm. Because of the stability of the droplet generator, nanovial outer diameter remained consistent in size with coefficients of variation of 2.48% and 2.02% for the two types of nanovials respectively (Supplementary Figure 1). Varying capping particle sizes were also manufactured from 30 to 40 µm with CVs of 2.81% and 2.44%. Although capping of nanovials was achieved across a range of capping particle diameters spanning 25 µm to 90 µm, we used capping particles of 30 µm diameter to cap 40 µm nanovials and capping particles of 40 µm diameter to cap 70 µm nanovials. Nanovials and capping particles are functionalized with biotin using sulfo-N-hydroxysuccinimide-biotin (NHS-biotin) conjugation to free lysine residues on the entrapped gelatin, to enable downstream cell and molecular capture (Supplementary Figure 2).

We identified key process parameters and chemistries that maximized the capping particle-nanovial docking process and its stability. To assemble capped nanovials, nanovials are mixed with capping particles and induced to interact through centrifugation steps in a wash buffer (WB) composed of phosphate buffer saline and Pluronic F-127 or media. Although we could not visualize the capping process under centrifugation, we observed the mixed system of nanovials and capping particles settling under gravity to inform processes expected to occur during centrifugation, but at a slower rate (Video S1). A few insights from these observations were: (i) nanovials settle and orient with their cavities facing upwards due to their asymmetric center of mass, and (ii) less crosslinked and dense capping particles then settle into the cavities of nanovials, and (iii) because both nanovials and capping particles can rotate under force moments, capping particles that fall off-center into a nanovial cavity, appear to re-align the nanovial to form a capping event. Based on these observations we surmise that capping rates are high even with the stochastic nature of the random encounters between capping particles and nanovials, because the system has a broad range of starting conditions and trajectories that favor coupling. Reflecting the stochasticity of the process, increasing the ratio of capping particles to nanovials improves capping efficiency (Figure 2A). When mixing unfunctionalized capping particles of 40 μm diameter with nanovials of 70 μm outer diameter at a ratio of 5:1 we observed capping rates of 9.3% ± 5.4% as measured by flow cytometry (Figure 2A). Capping rates appeared to saturate around 10% for even higher ratios, such as 10:1. The ability to interact with open nanovials was enhanced by mixing and repeat centrifugation steps, with increasing rates of capping from 8.7% ± 4.1% to 18.0% ± 2.5% with subsequent centrifugations (Figure 2B). Centrifugation speed or time had minimal effects on formation of capped particles (Supplementary Fig. 3A). Similarly, the vessel size in which capping was performed did not significantly impact capped particle formation (Supplementary Fig. 3B). Although physical docking between the flexible hydrogels enables stable capping, molecular affinity also was found to further stabilize interacting particles. Streptavidin-biotin interactions on the complementary particles promoted capped nanovial formation. Interestingly, we found that streptavidin functionalization of capping particles interacting with biotinylated nanovials led to highest capping rates (17.7% ± 4.0%), outperforming streptavidin on nanovials interacting with biotinylated capping particles (11.9% ± 5.1%), streptavidin on both particle types (10.7% ± 3.1%) and no functionalization (4.7% ± 0.6%) (Figure 2C). For these experiments, a single centrifugation step was performed with a capping particle to nanovial ratio of 5:1. While we evaluated these various factors on capping performance for larger 70 µm nanovials, well suited for mammalian cell assays, we found that using smaller particles (40 μm nanovials and 30 μm capping particles) further improved capping efficiency (Figure 2D). When transitioning the assay from wash buffer used for optimization studies to yeast culture media (YM) we observed even further improved capping efficiency (40.7% ± 6.6%). Taken together these methods enable consistent capping efficiency and rapid generation of hundreds of thousands to millions of compartments.

**Figure 2.**
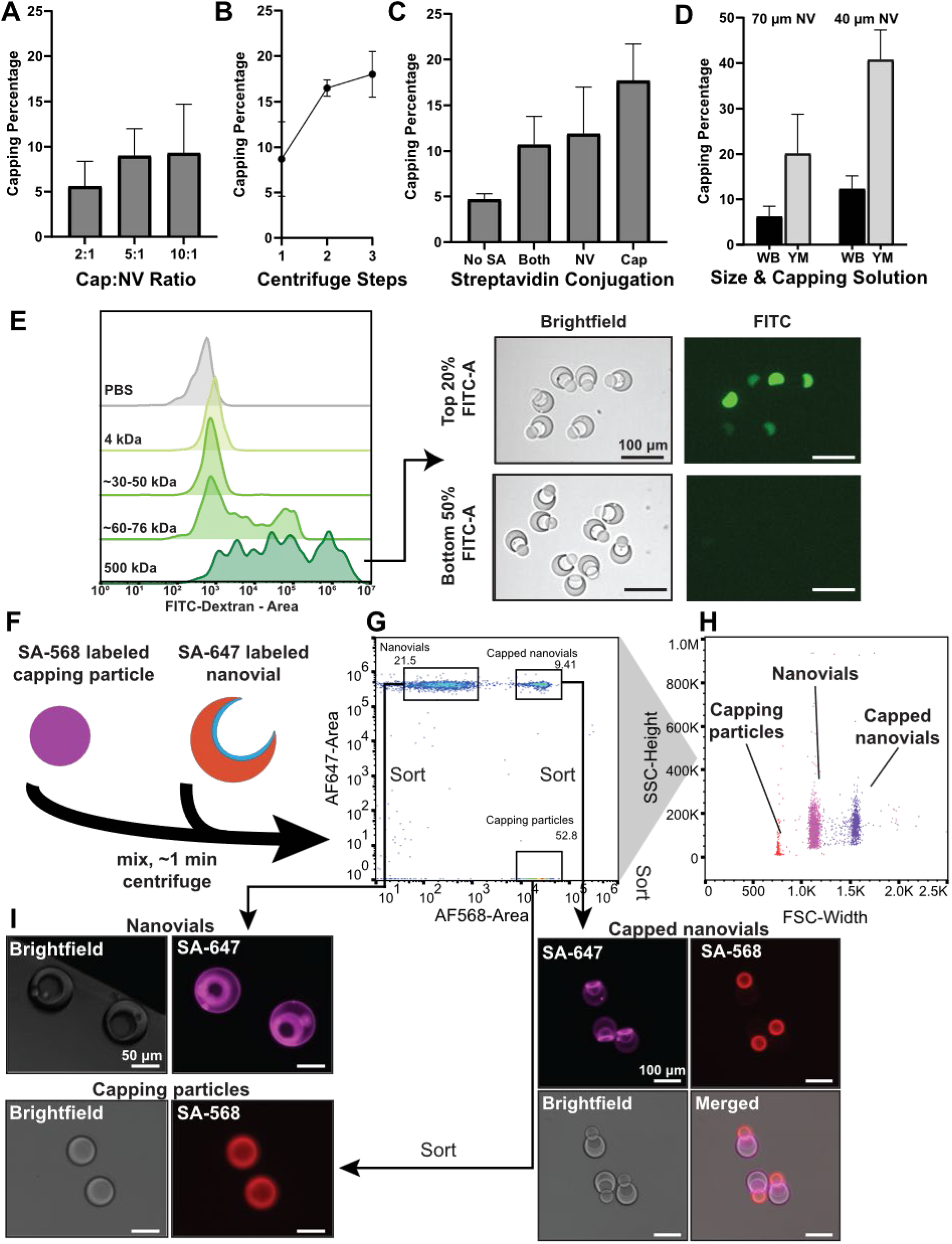
Capped nanovial formation characterization, molecular retention, and flow cytometry profiling. **A)** Effect of numerical capping particle to nanovial (NV) ratio on capping efficiency. **B)** Effect of multiple centrifugation steps on capping efficiency. **C)** Effect of streptavidin (SA) conjugation of nanovials and capping particles on capping efficiency. **D)** Effect of particle size (40 µm and 70 µm) and capping solution (WB – wash buffer, YM – yeast media) on capping efficiency. **E)** Flow cytometry and fluorescence microscopy showing retention of varying molecular weight FITC-dextran molecules. **F)** Hydrogel particles can be independently stained with fluorescently-bound streptavidin to barcode and detect capped nanovial events. **G)** Capped nanovials and constitutive particles can be identified on flow cytometry based on fluorescence. **H)** Each particle type has a unique forward and side scatter profile. **I)** Brightfield and fluorescence microscopy images of streptavidin-fluorophore-conjugated nanovials, capping particles, and capped nanovials following gating and sorting.

The docking of a capping particle onto the nanovial’s cavity opening forms a sealed microscale compartment that limits convective mixing and restricts the diffusion of larger molecules, while still permitting exchange of smaller nutrients and reagents. To characterize molecular transport across the capped nanovial system, we introduced fluorescent dextrans of varying molecular weights prior to capping and subsequently removed any unencapsulated dextran by washing. Dextrans smaller than 50 kDa readily diffused through the hydrogel matrix and/or the cap– nanovial interface (Figure 2E), indicating permeability to small solutes. In contrast, dextrans larger than 500 kDa were largely retained within the cavity, remaining confined for hours to days in the majority of capped nanovials. These results highlight the ability of capped nanovials to act as semi-permeable microcompartments, selectively isolating larger biomolecules while allowing controlled molecular exchange.

### Capped nanovial compatibility with standard flow sorter technology

Capped nanovials were compatible with analysis by flow cytometry and flow sorting without breakup of the composite asymmetric particle, similar to nanovials alone.^18^ We evaluated the ability to analyze and sort capped nanovials using standard FACS (Sony SH800S). We first functionalized both capping particles and nanovials with streptavidin Alexa Fluor 568 and streptavidin Alexa Fluor 647, respectively, after biotinylation (Figure 2F). Following docking, capped nanovials exhibited composite fluorescence profiles containing signal from both component particles, confirming successful assembly and maintenance of the docked form through fluid dynamic shear present during flow analysis (Figure 2G). This labeling allowed us to generate distinct fluorescent profiles for all three particle types (capping particles, nanovials, and capped nanovials) which could be gated to identify the unique forward scatter (FSC) and side scatter (SSC) profiles of each (Figure 2H). In particular, all of the three particle types had distinct FSC-width values (Figure 2H). Interestingly, the asymmetric capped nanovials did not yield a large distribution of scatter width profiles as would be expected for random orientations in the flow stream. Notably, the distinct FSC width value measured was larger than either the nanovials or capping particles alone. These results suggest that capped nanovials may align under the hydrodynamic sheath flow of the Sony SH800S microfluidic sheath flow chip, such that the long axis of the oblong particle is directed along the flow direction.

We further established that capped nanovials could be sorted without disrupting the capping particle-nanovial docking. Because of the larger particle sizes compared to single cells and nanovials alone the drop delay increment, which modulates the time between detection and the electrostatic deflection of a sorted droplet, was modulated to achieve maximum sorting performance. When using a Sony SH800S sorter, sort efficiency and purity metrics increase to a maximum using the Single Cell sort mode as the drop delay is adjusted, respectively (Supplementary Figure 4). A maximum sort yield of ∼50% and sort purity of ∼80% of 40 µm capped nanovials were achieved at the optimal drop delay, which approaches expected sort yields for other large particles.^18^ Reduced sorting yield was observed for larger capped nanovials with a shifted optimal drop delay. Sorted capped nanovials, gated on FSC-width, remained intact through hydrodynamic focusing, droplet-in-air sorting and recovery into a well plate demonstrating the robustness and stability of capping (Figure 2I). When gating based on FSC-width other particle populations could also be sorted and successfully identified (Figure 2I). Together, these results demonstrate that capped nanovials can be directly integrated into FACS-based pipelines, allowing for enrichment and recovery for downstream assays and analysis.

### Capped nanovials enable single-cell growth and selective enrichment

Capped nanovials compartmentalize growing colonies of cells supporting longitudinal tracking and enrichment based on growth rate. We first performed a growth assay using HyHEL5 hybridomas encapsulated within capped nanovials. Following nanovial functionalization with biotinylated anti-mouse CD45, cells were counted, loaded into nanovials based on binding to surface expressed CD45, subsequently isolated through capping and incubated for growth (Figure 3A). Calcein AM (622 Da) is permeable to capped nanovials, transporting through the hydrogel and/or interface between the capping particles and nanovials to stain cells within. Flow cytometry analysis revealed a distinct subpopulation of cell-loaded, capped nanovials, identifiable by high calcein AM fluorescence and high FSC-width (Figure 3A). When gating this population, sorting, and imaging it, the majority of events comprised capped nanovials containing cells. Capped nanovials with high calcein AM signal were sorted and cultured for 96 hours to monitor clonal expansion. Within this time, HyHEL5 cells proliferated within the compartment, forming visible colonies that were retained (Figure 3B). By the 96-hour time point, the mechanical force of the proliferating cells led to the displacement of the capping particle from some nanovials, allowing the colony to erupt from the compartment (Figure 3B) and colonize the well. This enables downstream recovery and facile recovery of cells compared to chemical de-emulsification steps required for microdroplets, as cells simply grow out of the compartment and expand into a standard culture vessel. These results indicate that capped nanovials accommodate several rounds of mammalian cell division while maintaining physical containment.

**Figure 3.**
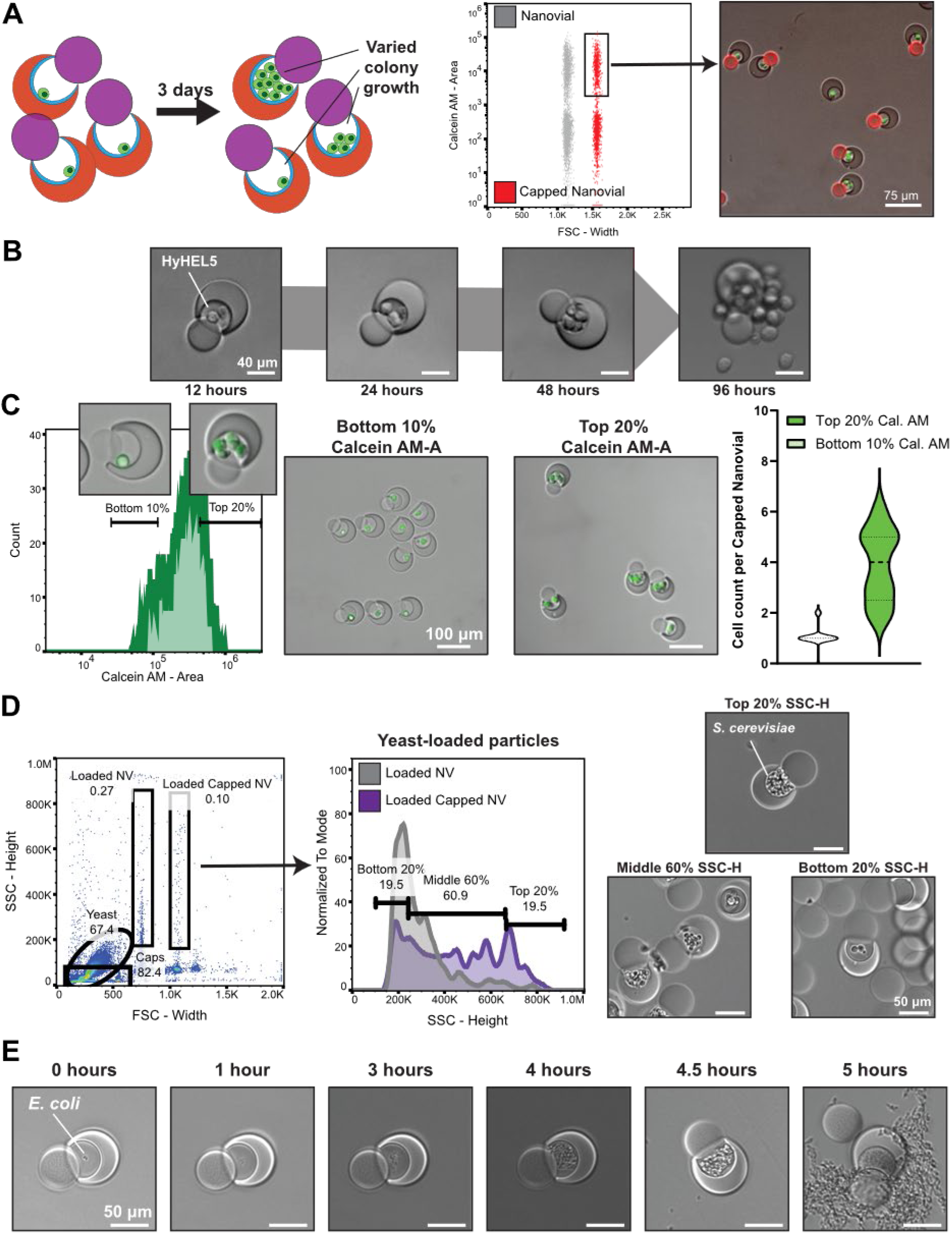
Capped nanovial growth assay. **A)** Flow cytometry profile of cell-loaded, capped nanovials. Forward scatter width and high calcein AM area signals are used to identify HyHEL5-loaded capped nanovials. **B)** Following sorting, HyHEL5 colonies proliferate within the particle compartment. After 96 hours the cell colony erupts from the enclosure. **C)** Following 72 hours of growth, HyHEL5-loaded capped nanovials are gated for Calcein AM area signal from flow cytometry and sorted. The top 20% of samples have on average four cells per compartment and the bottom 10% have on average one cell per compartment. **D)** Gating strategy for enrichment of yeast-loaded capped nanovials based on SSC height signal and images of *S. cerevisiae* colonies gated based on SSC post-sort. **E)** Timelapse images of *E. coli* growth over time in capped nanovials.

We selected colonies of hybridoma based on relative growth rate using the capped nanovials to enrich highly proliferative single cells. HyHEL5 cells were again loaded into nanovials and capped followed by 72 hours of culture to limit eruption from the capped nanovials. After staining with calcein AM, we compared the average number of cells per compartment following gating and sorting based on different thresholds of fluorescence intensity. Nanovials in the top 20% of the calcein AM signal distribution contained, on average, four cells per compartment, while those in the bottom 10% averaged approximately one cell per compartment (Figure 3C).

Using similar approaches, we also were able to isolate and sort populations of yeast and bacteria cells with varying growth rates in 40 µm nanovials with 30 µm capping particles. *S. cerevisiae* were loaded into nanovials via sealing with capping particles, cultured for 16 hours, sorted and selected based on growth using SSC-height signal, which reflected the density of the growing culture within the capped nanovial (Video S2, Figure 3D). Notably, compartmentalizing *S. cerevisiae* within capped nanovials enables formation of larger colonies than in nanovials alone by physically entrapping the cells (Figure 3D, middle panel). In another example, *E. coli* were loaded directly into the cavity of nanovials without any capture antibody, capped, and incubated over 5 hours, at which point the bacteria displaced the cap sufficiently to break containment (Video S3, Figure 3E). We established that *E. coli* expressing GFP loaded into capped nanovials for varying incubation periods could be identified and FACS-sorted based on overall GFP fluorescence of a colony confirmed by microscopy (Supplementary Figure 5A-B). These results demonstrate the ability to identify and isolate single-cell-derived colonies of mammalian cells, yeast, and bacteria based on growth from a mixed population in a high-throughput manner without the need for microfluidic systems at the time of the assay.

### Conducting single-cell secretion assays in capped nanovials reduces crosstalk

The enclosed cavity of capped nanovials confined secreted proteins within each local compartment improving the signal-to-noise ratio (SNR) in single-cell antibody secretion assays. COMSOL simulations of hybridomas loaded into nanovials generated higher local antibody concentrations within capped nanovials compared to uncapped nanovials over the same secretion period (Supplementary Figure 6). The concentration of secretions increased by 3.1-fold within the nanovial cavity compared to the uncapped nanovial when integrating over the cavity spaces of both particles. HyHEL5 hybridomas, which secrete mouse IgG antibodies against hen egg lysozyme (HEL), were loaded into both capped and uncapped nanovials functionalized with anti-mouse CD45 antibodies. In the capped configuration, biotinylated HEL antigen was conjugated to the spherical capping particle, while in the uncapped control it was bound to the nanovial surface along with anti-CD45 (Figure 4A). Cells were loaded into both systems for 90 minutes to enable adhesion and capped nanovials were formed. A fraction of the nanovials remained uncapped, serving as controls. IgG-specific signal on nanovials and on the capping particles docked onto nanovials both were on average elevated for flow cytometry analyzed events containing HyHEL5 cells compared to background nanovials or capped nanovials without cells (Figure 4B). However, the capped nanovial configuration exhibited an increased fold change enabling better discrimination between HyHEL5-loaded and empty capped nanovials (SNR = 5.59) compared to uncapped nanovials (SNR = 1.44) (Figure 4C). The interlocking architecture likely limits convective mixing and transport of secreted antibodies, leading to increased local concentration and reduced signal accumulation in neighboring compartments. The system still remains permeable to fluorescent labeling antibodies during mixing/staining steps. Since capping particles containing binding antigen are introduced following cell loading, crosstalk introduced during loading steps that occur for standard nanovials may also be reduced. Following sorting capped nanovials or uncapped nanovials based on high fluorescence intensity signal for secretion specific signal, fluorescence microscopy of capped nanovials reveals localized signal on caps associated with secreting cells, while crosstalk is observed directly on nanovials without bound cells (Figure 4C). Fluorescent signal localized near single cells exclusively within the capped nanovial indicated efficient and spatially resolved capture of secreted antibodies. We also found that when gelatin-containing biotinylated-HEL-conjugated capping particles and nanovials were incubated with varying dilutions of HyHEL5 supernatant, capping particles exhibited a slightly larger dynamic range across dilutions from 60 to 4250 intensity values, compared with 200 to 3100 intensity values (Supplementary Figure 7) and this extended range of intensity on capping particles enabled resolution of different secretion amounts accumulating in a time-dependent manner over an order of magnitude in intensity when incubating up to 6 hours (Supplementary Figure 8). Together, these results highlight the ability to limit crosstalk and introduce time dependent capture of secretions using capped nanovials enhancing the ability to select functional cells within a complex background.

**Figure 4.**
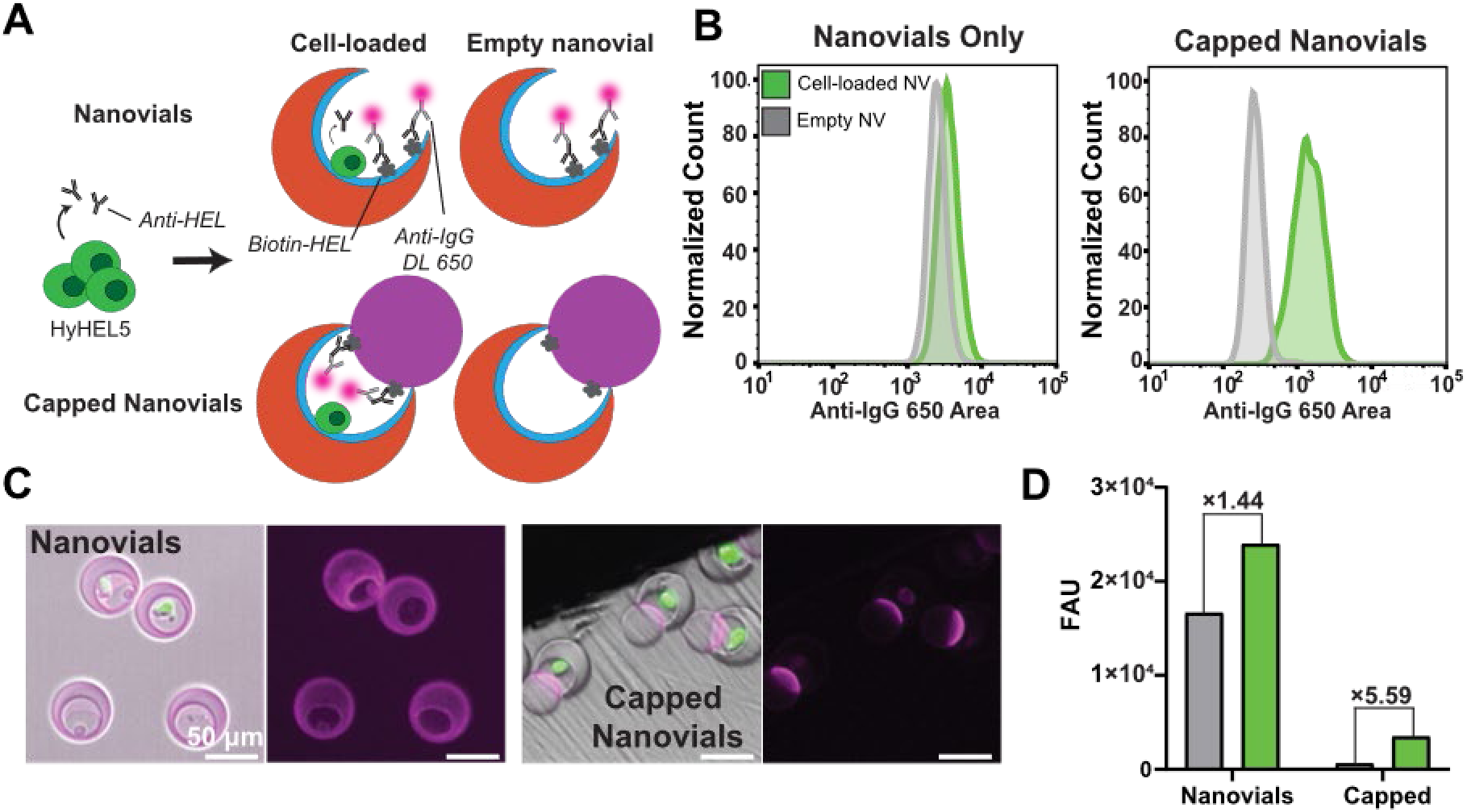
Capped nanovial antibody secretion assay. **A)** HyHEL5 hybridomas are loaded into capped and uncapped nanovials with biotinylated HEL protein conjugated to the capping particle and the nanovial, respectively. Secreted antibodies are detected with fluorescent anti-mouse IgG. **B)** Flow cytometry histogram comparisons of fluorescence secretion signal for cell-loaded and unloaded nanovials and cell-loaded and unloaded capped nanovials. **C)** Fluorescence and brightfield microscopy images of the single-cell secretion assay results for ab assay on nanovials and capped nanovials. **D)** MFI of signal for unloaded nanovials and capped nanovials (grey) and cell-loaded uncapped and capped nanovials (green). Capped nanovial workflow led to an increased signal-to-noise ratio compared to nanovials alone.

### Measuring and sorting cells based on cell-cell interactions within capped nanovials

We hypothesized that we could compartmentalize and maintain interacting cells within capped nanovials to accumulate secreted molecules and amplify their interactions. Specifically, we co-loaded OKT3 hybridomas (which constitutively secrete anti-CD3 antibodies) with NFAT-GFP Jurkat reporter cells, which express GFP upon T cell activation via CD3 binding and clustering (Figure 5A). When both cells are confined within a single capped nanovial, the local accumulation of anti-CD3 secreted by the hybridoma was expected to activate the Jurkat cell, triggering GFP expression. Following incubation for 24 hours we observed a marked shift in the GFP fluorescence in capped nanovials containing OKT3 cells compared to capped nanovials with only Jurkat reporter cells by flow cytometry (Figure 5B). Flow cytometry analysis confirmed that reporter cell activation was robust and distinguishable in capped nanovials (12.6-fold average signal increase between OKT3+/OKT3-containing capped nanovials, n = 3 biological replicates using different cultures of both OKT3 and Jurkat reporter cells) (Figure 5C). Fluorescence microscopy of sorted capped nanovials with high GFP levels confirmed successful co-encapsulation of Jurkat and hybridoma pairs and generation of GFP fluorescence in the activated reporter cells when co-encapsulated (Figure 5D).

**Figure 5.**
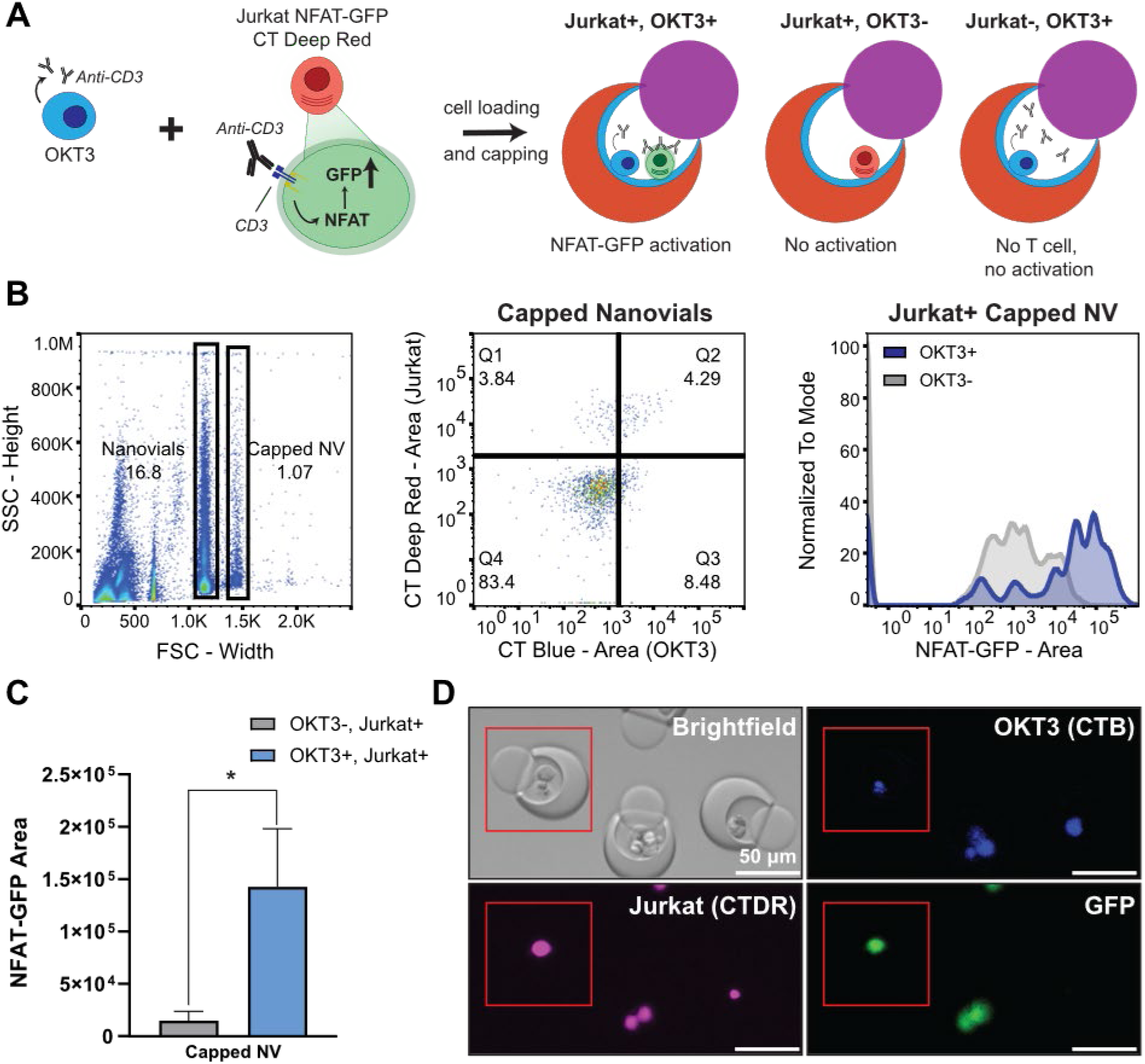
Two-cell interaction assay. **A)** Schematic of co-loading OKT3 hybridomas (CellTracker Blue) with Jurkat NFAT-GFP cells (CellTracker Deep Red) into capped nanovials. **B)** Gating strategy to identify Jurkat-loaded capped nanovials with OKT3 hybridoma (Q2) and without OKT3 hybridoma (Q1). NFAT-GFP area profiles for Jurkat+ capped nanovials with (Blue histogram, Q2) and without (Grey histogram, Q1) OKT3 cells shows distinct activation profiles. **C)** Mean fluorescence intensity of events in Q2 versus Q1 shows a significant increase, * = p < 0.05, n = 3 replicates. **D)** Fluorescence microscopy of sorted OKT3 positive and Jurkat positive capped nanovials (Q2).

To assess the utility of this platform to identify functional antibody clones in a background of non-functional antibodies we performed a population spiking experiment using an 80% background of HyHEL5 hybridomas (non-stimulatory) with a 20% population of OKT3 cells (Figure 6A). After co-loading the mixed hybridomas with Jurkat NFAT-GFP cells, we again incubated for 24 hours to induce CD3-mediated activation and GFP expression. Following gating for Jurkat-loaded (CellTracker Deep Red positive) capped nanovials, we found an over 30-fold increase in the average GFP activation signal for capped nanovials co-loaded with OKT3 cells (CellTracker Blue positive) compared to HyHEL5 cells (CellTracker Orange) (Figure 6B), resulting in clear thresholds for selection between the two populations. Reflecting a blinded selection experiment we also gated co-loaded capped and uncapped nanovials for the top NFAT-GFP area signal, and identified the percentage of particles that were loaded with on-target OKT3 cells. We found that purity up to 100% for OKT3 hybridoma could be achieved for the most stringent gates, and across every NFAT-GFP area gating stringency capped nanovials yielded improved purity compared to uncapped controls (Figure 6C). Fluorescence microscopy images of representative co-loaded capped nanovials with OKT3 or HyHEL5 hybridoma reflected the selective induction of GFP expression in the reporter cells co-loaded with OKT3 (Figure 6D).

**Figure 6.**
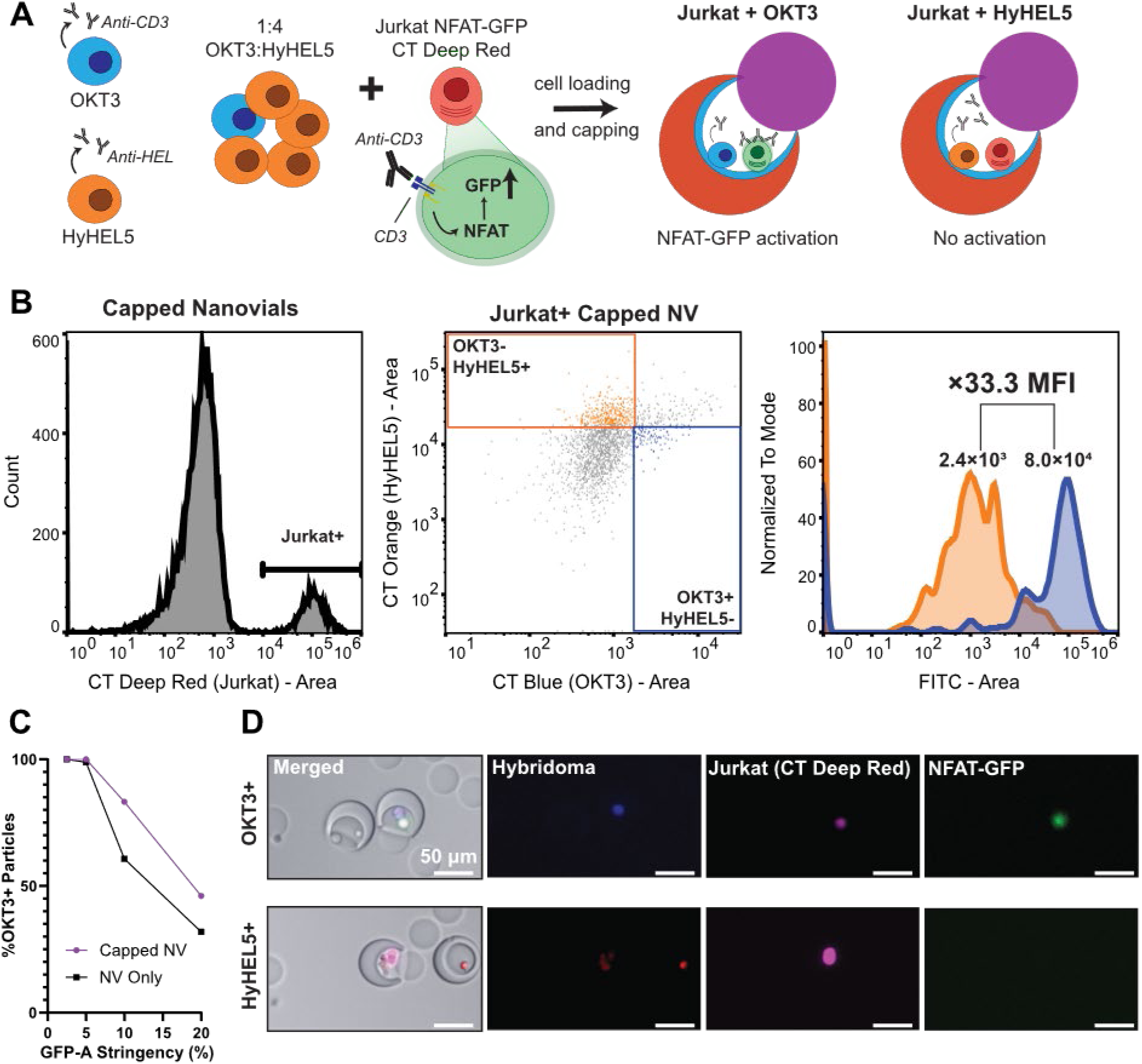
Two-cell spiked interaction assay in capped nanovials. **A)** Schematic of hybridoma spiking and co-loading with Jurkat NFAT-GFP cell line into capped nanovials. **B)** Flow cytometry analysis of capped nanovials loaded with Jurkat cells and multiple hybridoma populations. After gating for Jurkat reporter cell presence via CellTracker Deep Red, events are gated for single positive hybridoma populations and histograms of NFAT-GFP activation fluorescence are shown for OKT3+HyHEL5-(blue) and OKT3-HyHEL5+ events (orange). A >30-fold increase in GFP MFI is observed for on-target OKT3 hybridoma co-loaded with Jurkat reporter cells. **C)** Comparison in OKT3 enrichment when gating based on NFAT-GFP signal area alone at different stringencies for capped nanovials and uncapped nanovials. **D)** Fluorescent microscopy images of representative capped nanovials containing a single Jurkat NFAT-GFP reporter cell and either one OKT3 or HyHEL5 hybridoma.

## Discussion

The capped nanovial platform introduced here redefines the functionality of lab-on-a-particle systems by enabling stable compartmentalization, multi-cell assays, and growth-based selections—all within an accessible, scalable format. By simply combining two hydrogel particles through complementary docking, we create sealed, suspendable vessels that bridge a critical gap in microscale biological experimentation.

A major advance of the capped nanovial system is its ability to support cell growth assays within a confined microenvironment. While prior nanovial platforms enabled secretion assays, their open architecture was less suited for capturing clonal expansion or measuring growth phenotypes. Here, the addition of a hydrogel capping particle transforms the open nanovial into a true microvessel— retaining single cells or small colonies over time, restricting exchange of larger molecules, and preserving secretions. This is, to our knowledge, the first demonstration of bacterial, yeast, and mammalian cell growth within nanovials, opening powerful new applications in directed evolution, synthetic biology, and biomanufacturing optimization where growth rates are critical readouts.

Beyond growth, the capped nanovials enable a new class of multi-cell assays. By co-encapsulating different cell types and sealing them together, secreted molecules can accumulate locally, amplifying functional interactions that would otherwise be diluted in bulk. We demonstrated this by co-loading antibody-secreting hybridomas with NFAT-GFP reporter T cells, achieving robust, selective activation dependent on local antibody production. This is the first time a reporter-based multi-cell assay has been implemented using nanovials, aided by the secure microenvironment established through capping.

The capped architecture also introduces an important design flexibility: the nanovial and capping particle can be independently functionalized with different capture agents, biomolecules, or barcodes. This modularity extends the range of possible assays from single-cell secretion analysis to antibody, TCR, and CAR discovery workflows, cell-cell interaction screening, and even colony or single-cell sequencing when capping particles are oligo-barcoded. In principle, capped nanovials could enable combined workflows where functional screening is followed by sequencing to link function to genotype at the level of a growing colony, a cell-cell interaction pair, or a single cell.

Perhaps most remarkably, the simplicity of the capped nanovial platform stands out. While much of microfluidics and single-cell analysis relies on specialized instrumentation and expertise, capped nanovials are assembled and analyzed with standard laboratory tools—simple mixing, centrifugation, and flow cytometry. This operational simplicity, coupled with the robustness of the capped compartments even through sorting, positions the platform for broad adoption across biology, biotechnology, and medicine. Taken together, the capped nanovial system provides a new approach for performing scalable, accessible assays of growth, secretion, and interaction at the single-cell and colony level. By miniaturizing and democratizing the concept of a sealed experimental vessel, this platform extends the power of high-throughput biology—bringing the classic virtues of the test tube and the petri dish into the microscale era, and supercharging collection of functional cell data to power the biological AI models of the future.

## Materials and methods

### Nanovial particle fabrication

Nanovial production has been described in detail in previous publications.^13,14^ Nanovials were fabricated using a polydimethylsiloxane (PDMS) microfluidic device designed with a flow-focusing geometry for droplet generation. The aqueous polyethylene glycol (PEG) phase consisted of 27.5% (w/v) 4-arm 5 kDa PEG-acrylate (Advanced BioChemicals, 4AP0902-1g) and 4% (w/v) lithium phenyl-2,4,6-trimethylbenzoylphosphinate (LAP) (Sigma, 900889), dissolved in phosphate-buffered saline (PBS) (Thermo Fisher, 14190250). This was co-injected with a separate aqueous gelatin phase containing 20% (w/v) cold-water fish gelatin (Sigma, G7041100G) in sterile-filtered deionized water. An oil phase composed of 1% (w/w) 008-FluoroSurfactant (RAN Biotechnologies, 008-FluoroSurfactant-1G) in Novec™ 7500 fluorinated oil (3M, 7100134816) served as the continuous phase. Each solution was loaded into individual syringes and introduced into the device via syringe pumps at flow rates of 1 μL min^-1^ (PEG), 1 μL min^-1^ (gelatin), and 15 μL min^-1^ (oil). Upon reaching a stable flow regime, monodisperse droplets were formed with spontaneous phase separation between the PEG-rich and gelatin-rich domains. Crosslinking of the PEG phase was initiated by UV exposure through a 10× microscope objective. Following droplet formation and photo-crosslinking, the emulsion was collected into a microcentrifuge tube. Residual surfactant was removed by triple washing with sterile-filtered Novec™ 7500. Droplets were then demulsified by adding a 20% (v/v) solution of Pico-Break (Sphere Fluidics, C082) in Novec™ 7500. Oil was aspirated, and remaining traces were eliminated through three hexane washes. Excess hexane was subsequently removed via aspiration, and particles were further purified by washing in sterile-filtered 70% (v/v) ethanol three times. To eliminate aggregates, the nanovials were passed through a 70 μm cell strainer (Stem Cell Technologies, 27216), then incubated overnight at 4 °C in ethanol for sterilization. Nanovials were then incubated overnight at 4 °C in a solution of 10 mM biotin-NHS (ApexBio, A8001). Prior to use, particles were washed in a Pluronic-based buffer composed of 0.05% (w/v) Pluronic F-127 (Sigma, P2443), 1% (v/v) antibiotic-antimycotic, and 0.5% (w/v) bovine serum albumin (BSA) (Sigma, A7906). Nanovials were stored at 4 °C in this buffer until use.

### Capping particle fabrication

Capping particle production was performed similarly to nanovial fabrication. Capping particles were fabricated using a flow-focusing PDMS device with a 28 µm junction height to create 30 and 40 µm particles. The aqueous PEG phase used 12% (w/v) 4-arm 5 kDa PEG-acrylate and 4% LAP dissolved in PBS. The second aqueous phase used a 2% (w/v) solution of gelatin dissolved in deionized water. An oil phase composed of 0.5% (w/v) of Pico-Surf surfactant (Sphere Fluidics, C024) in Novec™ 7500 fluorinated oil was used as the continuous phase. The PEG and gelatin phases were mixed in a 1:1 volumetric ratio and loaded into two syringes with the oil phase in a third syringe. The solutions were perfused through the flow-focuser at flow rates of 0.5 μL min^-1^ (PEG + gelatin), 0.5 μL min^-1^ (PEG + gelatin), and 18 μL min^-1^ (oil) for 30 µm particles and at 1 μL min^-1^ (PEG + gelatin), 1 μL min^-1^ (PEG + gelatin), and 15 μL min^-1^ (oil) for 40 µm particles. Upon reaching a stable flow regime, monodisperse droplets were formed with a uniform distribution of PEG and gelatin throughout each droplet. Crosslinking was performed through UV exposure through a 10× microscope objective, which polymerized the PEG and embedded gelatin strands within the matrix. Following particle collection, residual surfactant was removed by triple washing with sterile-filtered Novec™ 7500. Droplets were then demulsified by adding a 20% (v/v) solution of perfluoro-1-octanol (PFO) (Sigma, 370533) in Novec™ 7500. Oil was aspirated, and remaining traces were eliminated through three hexane washes. Capping particles were then sterilized, filtered, biotinylated, and transferred to a Pluronic-based buffer similarly to nanovials.

### Capped nanovial formation

1.8 × 10^5^ 70 μm nanovials were incubated with 50 μL of 300 μg mL^-1^ streptavidin (Thermo Fisher, 434302) in a 1.5 mL microcentrifuge tube. In a separate 1.5 mL microcentrifuge tube, 9.0 × 10^5^ 40 μm capping particles were incubated with 75 μL of 300 μg mL^-1^ streptavidin. Both solutions are incubated for 30 minutes at room temperature, then washed three times with Pluronic washing buffer. Both particle types are then mixed together in a 0.5 mL Eppendorf tube and resuspended to a final volume of 300 μL in Pluronic washing buffer. The tube containing capping particles and nanovials is then centrifuged at 600 RCF for 90 seconds, creating a mixed population of capped nanovials, free nanovials, and free capping particles. For testing the optimal condition for forming capped nanovials, a variety of different conditions were tested: the numeric ratio of capping particles to nanovials (2:1, 5:1, and 10:1), frequency of centrifugation steps (1∼3), different combinations of capping particles and nanovials with or without 300 μg mL^-1^ streptavidin functionalization, varying the relative centrifugal force (RCF) of the centrifugation step (150 RCF∼ 1400 RCF), centrifugation duration (30 sec ∼ 120 sec), and the volume of vessel to perform capping formation (0.5 mL, 1 mL, and 5 mL). All flow cytometric analysis was performed using a Sony Biotechnology SH800S Cell Sorter. Alexa Fluor 588 streptavidin fluorophore was excited using a 561 nm laser filtered through a 617/30 filter. Alexa Fluor 647 streptavidin fluorophore was excited using a 638 nm laser filtered through a 665/30 filter. Sorting was performed on single-cell sort mode with a drop delay set to 15 (from a machine-calibrated setting of 18). Capping percentages were defined as capped nanovial events / (nanovial events + capped nanovial events), where all events were identified using the distinct forward scatter profile of the different particle populations. Capped nanovial sort efficiency and purity metrics were calculated by sorting 100 capped nanovial events based on FSC width and SSC height profiles and counting the ratio of sorted capped nanovials to desired number of capped nanovials (sort efficiency) and the ratio of sorted capped nanovials to all sorted particles (sort purity) (Supplementary Figure 1).

### Culture of hybridoma cells and reporter T cells

OKT3 hybridomas were purchased from ATCC. HyHEL5 hybridomas were provided by Richard Wilson from the University of Houston Department of Biology and Biochemistry. Jurkat NFAT-GFP cells were provided by Zhiyuan Mao and Owen Witte from the University of California Los Angeles Department of Microbiology, Immunology, and Molecular Genetics. Hybridoma cells were cultured in IMDM cell culture media (Thermo Fisher 12440053) supplemented with 10% fetal bovine serum (Thermo Fisher A5669701) and 1% antibiotic-antimycotic (Thermo Fisher 15240062). Jurkat cells were cultured in RPMI 1640 cell culture media (Thermo Fisher 11875093). Media was sterile filtered using 0.22 µm, 500 mL Stericups (Thermo Fisher S2GPU05RE). All cells were quickly thawed from liquid nitrogen at 1 × 10^5^ cells per mL and passaged three times per week. Cells were maintained in a sterile incubator at 37 °C and 5% CO_2_ and passaged up to P20 before replacement with a fresh vial. All cells were assessed for >95% viability before proceeding with experiments.

### Hybridoma growth assay

1.8 × 10^5^ 70 μm nanovials were incubated with 50 μL of 300 μg mL^-1^ streptavidin in a 1.5 mL microcentrifuge tube for 30 minutes at room temperature, then washed three times with Pluronic washing buffer. Nanovials were then incubated with 50 μL of 50 μg mL^-1^ biotinylated anti-mouse CD45 antibody (Thermo Fisher, 13045182). In parallel, 9.0 × 10^5^ 40 μm capping particles were incubated with 75 μL of 300 μg mL^-1^ streptavidin for 30 minutes at room temperature. Both particles were washed three times with Pluronic washing buffer. 3.0 × 10^5^ HyHEL5 cells were mixed in with the antibody-conjugated nanovials in the well of a 24 well plate and placed on a rocker in an incubator at 37 °C incubator for 90 minutes. Every 30 minutes the cells and nanovials were mixed by pipetting to promote cell loading. Following loading, unbound cells were removed using a 37 μm cell strainer (STEMCELL Technologies, 27215) and nanovials with and without cells were reverse strained into a 5 mL Eppendorf tube. This mixture is briefly centrifuged at 100 rcf for 2 minutes and the supernatant is removed. The cell-loaded nanovials are then transferred to a 0.5 mL Eppendorf tube along with the capping particles. Cells are then sealed in capped nanovials by centrifuging the mixture at 300 rcf for 90 seconds. The now capped cell-loaded nanovials are transferred to the well of a 6 well plate that has been pre-filled with 5 mL of IMDM media. After three days of culture, the cell-loaded particles are stained with 1 μg mL^-1^ of calcein AM (Thermo Fisher, C3099) for 30 minutes. After washing two times with Pluronic washing buffer, capped nanovials are transferred to FACS tubes for flow cytometry. Calcein AM was excited using a 488 nm laser filtered through a 525/50 filter. After gating for capped nanovials using FSC-width, capped nanovials loaded with Calcein AM-stained cells were gated using Calcein AM-area signal and the bottom 10% and top 20% segments of this population were sorted.

### *E. coli* culture, capped nanovial loading, and analysis

*Escherichia coli* strains were cultured in Difco™ LB (Luria-Bertani) broth, Miller (BD Biosciences) at 37°C with shaking speed of 250 rpm. Solid media culture plates were prepared with Difco™ LB Agar, Miller (BD Biosciences) in 10 cm petri dishes and incubated at 37°C. 40 µm nanovials and 30 µm capping particles were stored at concentrations of 2,500 particles/µL and 3,400 particles/µL, respectively, in 0.05% (w/v) Pluronic F-127 (Sigma-Aldrich) in PBS at 4°C. All experiments were conducted in 0.5 mL microcentrifuge tubes, and centrifugation steps were performed using a swing-bucket centrifuge. The general bacteria loading and capping protocol is as follows: an aliquot of nanovials was centrifuged at 630 RCF for 2 minutes to pellet the particles, and excess supernatant was carefully removed. Nanovials and bacteria were mixed in a 1:3 nanovial-to-bacteria ratio and gently mixed using a 1000 µL pipette. The nanovial-bacteria suspension was centrifuged at 630 RCF for 2 minutes to promote bacterial loading into the nanovial cavities. To enhance loading efficiency, the pipette mixing and centrifugation steps were repeated twice more, for a total of three cycles. Following bacterial loading, excess supernatant was removed, and caps were added at a 1:2 nanovial-to-cap ratio. The mixture was gently mixed by pipetting and centrifuged at 630 RCF for 2 minutes to facilitate capping. This capping process was repeated twice more, for a total of three cycles, to maximize capping efficiency.

### Culture and capping of *S. cerevisiae*

A strain of *S. cerevisiae* 651 with two plasmids HRPKS (-U) and NRPKS + ACPTE (-L) was used.^19^ Cells stored at -80°C in glycerol stock solution were thawed in 5 mL of yeast extract peptone dextrose (YPD) cell media and cultured at 25°C with shaking speed of 360 rpm (Amerex Instruments). After an overnight incubation, the cell suspension was diluted 1/100 prior to all capped nanovial experiments. For encapsulating single *S. cerevisiae* in capped particles, 200 µL of the diluted yeast cell suspension, 6.0 µL of 40 µm pelleted nanovials (180,000 nanovials) and 12.7 µL of 30 µm pelleted capping particles (900,000 capping particles) were added to a 0.5 mL microcentrifuge tube. Following a brief vortexing step (Vortex Genie 2), the suspension was centrifuged (Thermo Scientific, Legend Micro 21 Centrifuge) at 600 RCF for 60 seconds. The pellet was gently suspended in washing buffer, strained and washed with washing buffer using a 20 µm strainer (Sysmex, CellTrics 04-004-2325) to remove unbound cells, and then reverse strained with 2 mL of YPD media into a well of 6 well plate. Cells encapsulated in capped particles were incubated for 12 hours at 25°C without shaking prior to analysis and sorting with flow cytometry. After the incubation, the particles were collected and suspended in 400 µL of washing buffer and transferred to a flow tube for flow cytometry. Flow cytometry analysis and sorting was performed using a Sony Biotechnology SH800S Cell Sorter. The particle suspensions were gated for (1) capped nanovial events based on forward scatter (FSC) profile and (2) colony growth based on side scatter (SSC) profile, where elevated SSC profile corresponded to higher yeast biomass content within capped nanovials. Capped nanovials were sorted on single-cell sort mode with a drop delay set to 17 (from a machine-calibrated setting of 19).

### Hen egg lysozyme biotinylation

Recombinant hen egg lysozyme (HEL) (Aviva System Biology, OORA00201) was biotinylated using an EZ-Link^TM^ Sulfo-NHS-LC-biotinylation kit (Thermo Fisher, 21435) following the manufacturer’s instructions. After purification, concentration was measured using a spectrophotometer and the protein was stored at -20 °C.

### Hybridoma secretion and binding assay

2.5 × 10^5^ 70 μm nanovials were incubated with 75 μL of 300 μg mL^-1^ streptavidin in a 1.5 mL microcentrifuge tube for 30 minutes at room temperature, then washed three times with Pluronic washing buffer. Nanovials were then incubated with 75 μL of 50 μg mL^-1^ biotinylated anti-mouse CD45 antibody. In parallel, 1.25 × 10^6^ 40 μm capping particles were incubated with 125 μL of 300 μg mL^-1^ streptavidin for 30 minutes at room temperature. Following washing with Pluronic washing buffer, capping particles were incubated with 125 μL of 25 μg mL^-1^ biotin-HEL. All particles were washed with Pluronic washing buffer. 5.0 × 10^5^ HyHEL5 cells were loaded onto the nanovials as described previously and unbound cells were strained out. After transfer to a 0.5 mL Eppendorf tube and mixing with the biotin-HEL capping particles, the cell-loaded capped nanovials were sealed via centrifugation. The capped, cell-loaded nanovials were transferred to the well of a 6 well plate that was pre-filled with 5 mL of IMDM media and incubated for 30 minutes for secretion accumulation to occur. The particles were then stained with 125 μL of 50 μg mL^-1^ anti-mouse IgG DyLight 650 antibody (Abcam ab98715) and 1 μg mL^-1^ of calcein AM for 30 minutes in the dark. After a final wash with Pluronic washing buffer, cell-loaded capped nanovials were transferred to FACS tubes for flow cytometry analysis. The DyLight 650 fluorophore was excited using a 638 nm laser filtered through a 665/30 filter. After gating for cell loaded, capped nanovials as described previously, particles were plotted for DyLight 650 area signal.

### Reporter T cell activation assay (bulk)

2.5 × 10^5^ 70 μm nanovials were incubated with streptavidin as described previously. Nanovials were then incubated with 75 μL of 25 μg mL^-1^ biotinylated anti-mouse CD45 antibody and 25 μg mL^-1^ biotinylated anti-human CD45 antibody (BioLegend 368534) for 1 hour at room temperature and washed three times with Pluronic washing buffer. In parallel, 1.25 × 10^6^ 40 μm capping particles were incubated with streptavidin as described previously. 2.0 × 10^5^ OKT3 cells were labeled with 200 μM CellTracker Blue CMHC (Thermo Fisher C2111). 2.0 × 10^5^ Jurkat NFAT-GFP cells were labeled with 1 μM CellTracker Deep Red (Thermo Fisher C34565). After washing, cells were mixed together and loaded into nanovials as described previously. After straining, capping particles were added to nanovials via centrifugation in a 0.5 mL Eppendorf tube. Cell-loaded, capped nanovials were transferred to the well of a 6 well plate pre-loaded with 5 mL of a 50:50 volumetric mixture of complete IMDM and complete RPMI media and incubated for 24 hours to enable OKT3 antibody secretion and Jurkat reporter activation. Particles were transferred into Pluronic washing buffer and placed in FACS tubes for flow cytometry analysis. The CellTracker Blue CMHC fluorophore was excited using a 405 nm laser filtered through a 450/50 filter. The NFAT-GFP reporter fluorophore was excited using a 488 nm laser filtered through a 525/50 filter. The CellTracker Deep Red fluorophore was excited using a 638 nm laser filtered through a 665/30 filter. After gating for capped nanovials, events were gated for CellTracker Deep Red area and CellTracker Blue CMHC area signal. These events were then plotted for NFAT-GFP area signal to compare activation profiles of capped nanovials with and without hybridomas.

### Reporter T cell activation assay (spiked)

5.0 × 10^5^ 70 μm nanovials were incubated with 150 μL of 300 μg mL^-1^ streptavidin as described previously. Nanovials were then incubated with 150 μL of 25 μg mL^-1^ biotinylated anti-mouse CD45 antibody and 25 μg mL^-1^ biotinylated anti-human CD45 antibody for 1 hour at room temperature and washed three times with Pluronic washing buffer. 2.5 × 10^6^ 40 μm capping particles were incubated with 250 μL of 300 μg mL^-1^ streptavidin as described previously. 6.0 × 10^4^ OKT3 cells were stained with CellTracker Blue CMHC as described previously. 2.4 × 10^5^ HyHEL5 cells were stained with 20 μM CellTracker Orange CMRA. 5.0 × 10^5^ Jurkat NFAT-GFP cells were stained with CellTracker Deep Red as described previously. After washing excess cell stain, the different cells were mixed together and loaded into nanovials, strained, and capped as previously described. Following 24 hours of incubation for antibody secretion and Jurkat activation, particles were transferred to Pluronic washing buffer and placed in FACS tubes for flow cytometry. The CellTracker Orange CMRA fluorophore was excited with a 561 nm laser filtered through a 617/30 filter. After gating for capped nanovials, events were gated for the presence of Jurkat NFAT-GFP cells based on CellTracker Deep Red area signal. These events were then gated for CellTracker Orange CMRA area and CellTracker Blue CMHC signal. Events that were single positive for one of these markers were then plotted for NFAT-GFP area signal to compare the activation profiles of Jurkat-loaded capped nanovials with either type of hybridoma co-loaded.

### COMSOL simulation of IgG accumulation in capped nanovials

Secretion was modeled in a 2D axisymmetric COMSOL (Version 6.3) simulation using the Transport of Diluted Species module, assuming water as the solvent and purely diffusive transport. IgG was secreted at a constant rate of 1.71×10^-12^ mol m^-2^ s^-1^, equivalent to 1 pg hr^-1^ per cell, based on literature estimates for mouse hybridoma cells.^20^ The diffusion coefficient for IgG was set to 4×10^-11^ m^2^ s^-1^, consistent with reported values for IgG in water or PBS.^21^ The simulation was evaluated over 2 hours to generate IgG concentration profiles within each particle. The cap and the nanovial were simulated to not be permeable to the IgG species. In the capped nanovial simulation, a gap of 100 nm was used to simulate limited transport out of the compartment through the nanovial lip region.

## Data availability

The data supporting this article have been included as part of the electronic submission. Raw data and .FCS files are available upon request.

## Acknowledgments

Flow cytometry was performed in the UCLA Jonsson Comprehensive Cancer Center which is supported by National Institutes of Health awards P30 CA016042 and 5P30 AI028697. The work is partially supported by grant CA256084 from the National Institutes of Health and grant 2023332386 from the Chan Zuckerberg Initiative Donor Advised Fund (CZI DAF), an advised fund of the Silicon Valley Community Foundation. Y. N. is supported by postdoctoral fellowships from Japan Society for the Promotion of Science and Nakatani Foundation.

---

**Supplementary Figure 1.**
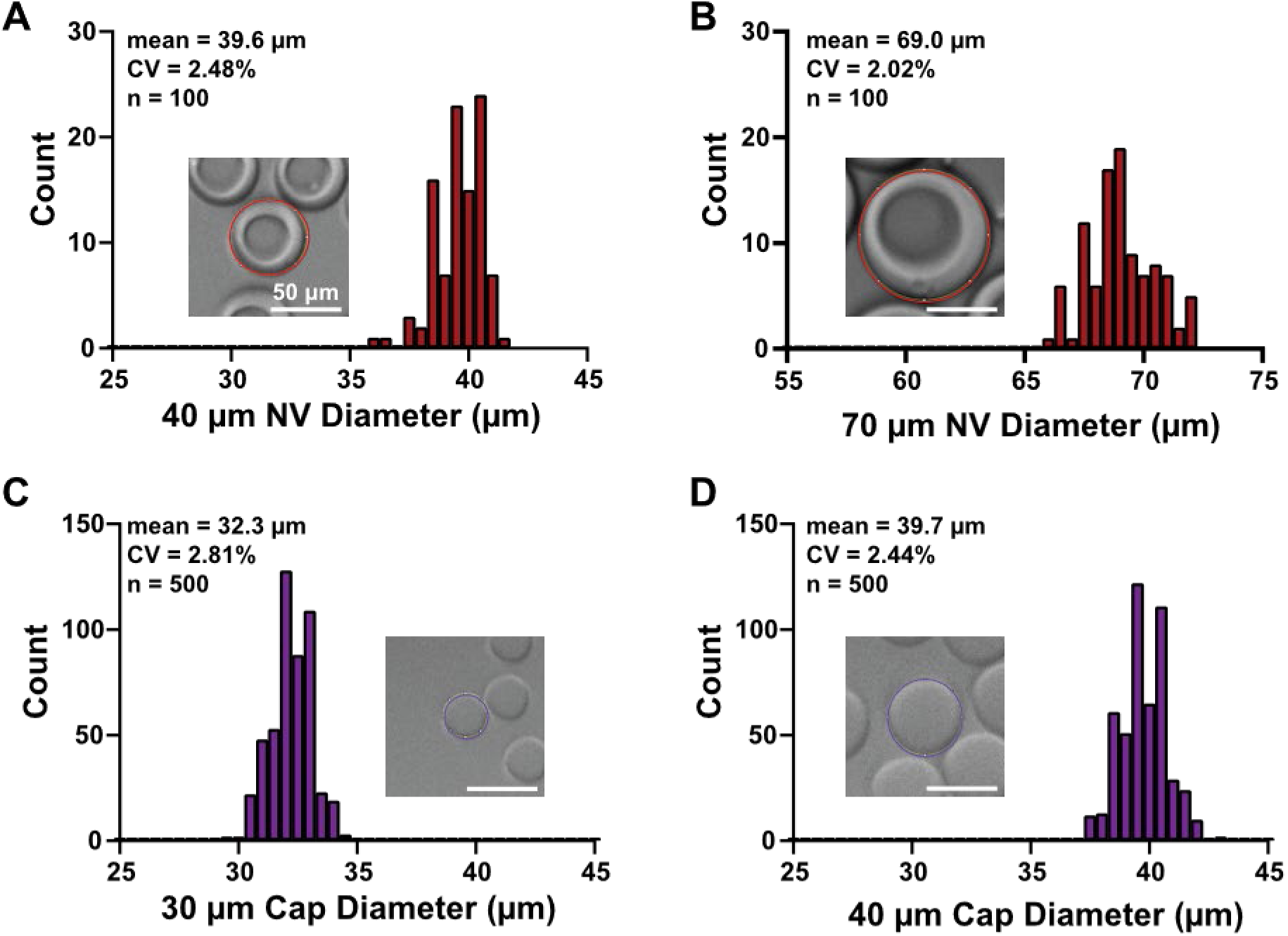
Capping particle and nanovial size measurements. Particle size and CVs of **A)** 40 µm nanovials, **B)** 70 µm nanovials, **C)** 30 µm capping particles, and **D)** 40 µm capping particles.

**Supplementary Figure 2.**
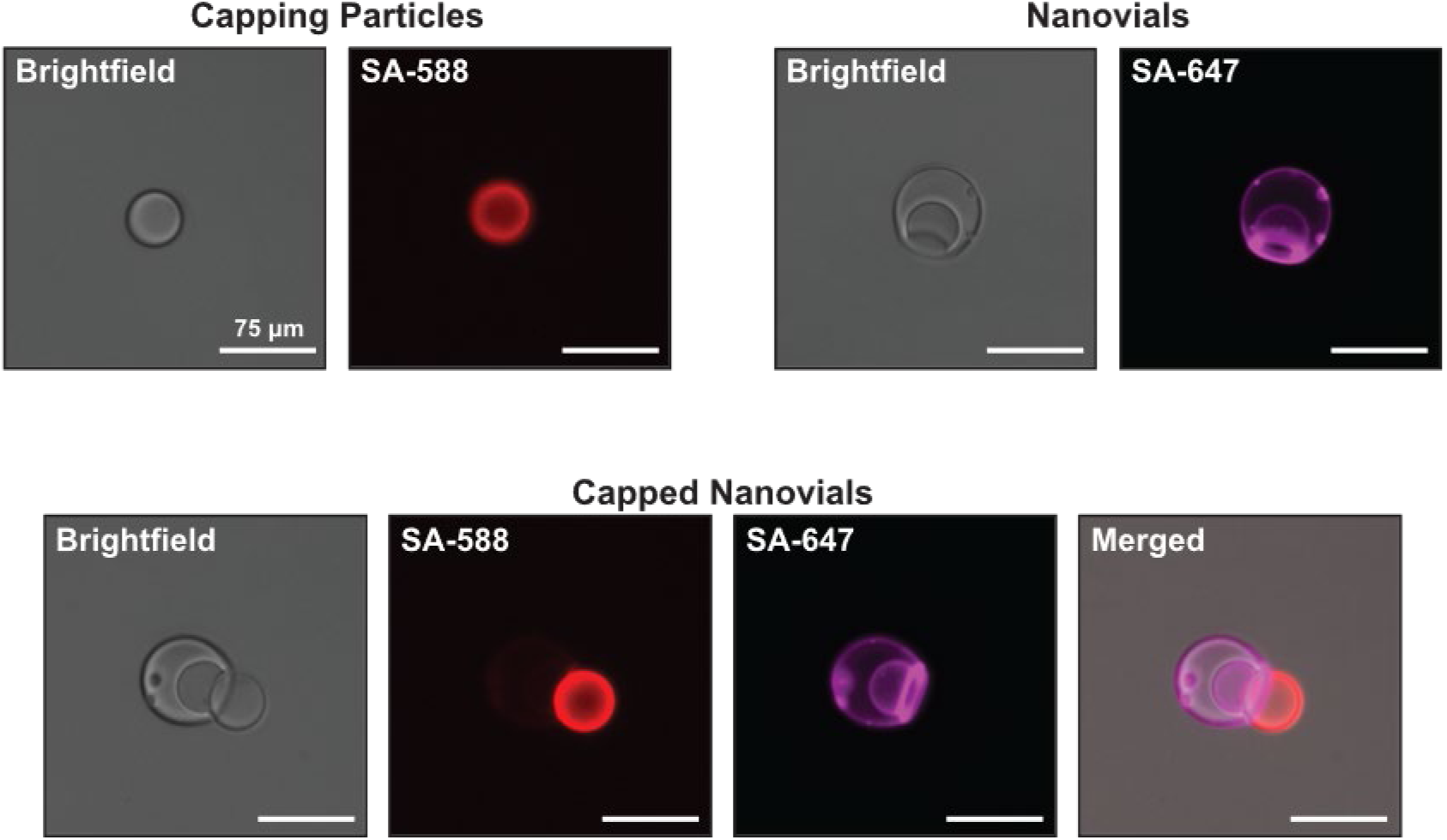
Biotin and fluorescent streptavidin conjugation to particles. Fluorescent streptavidin conjugation to capping particles and nanovials.

**Supplementary Figure 3.**
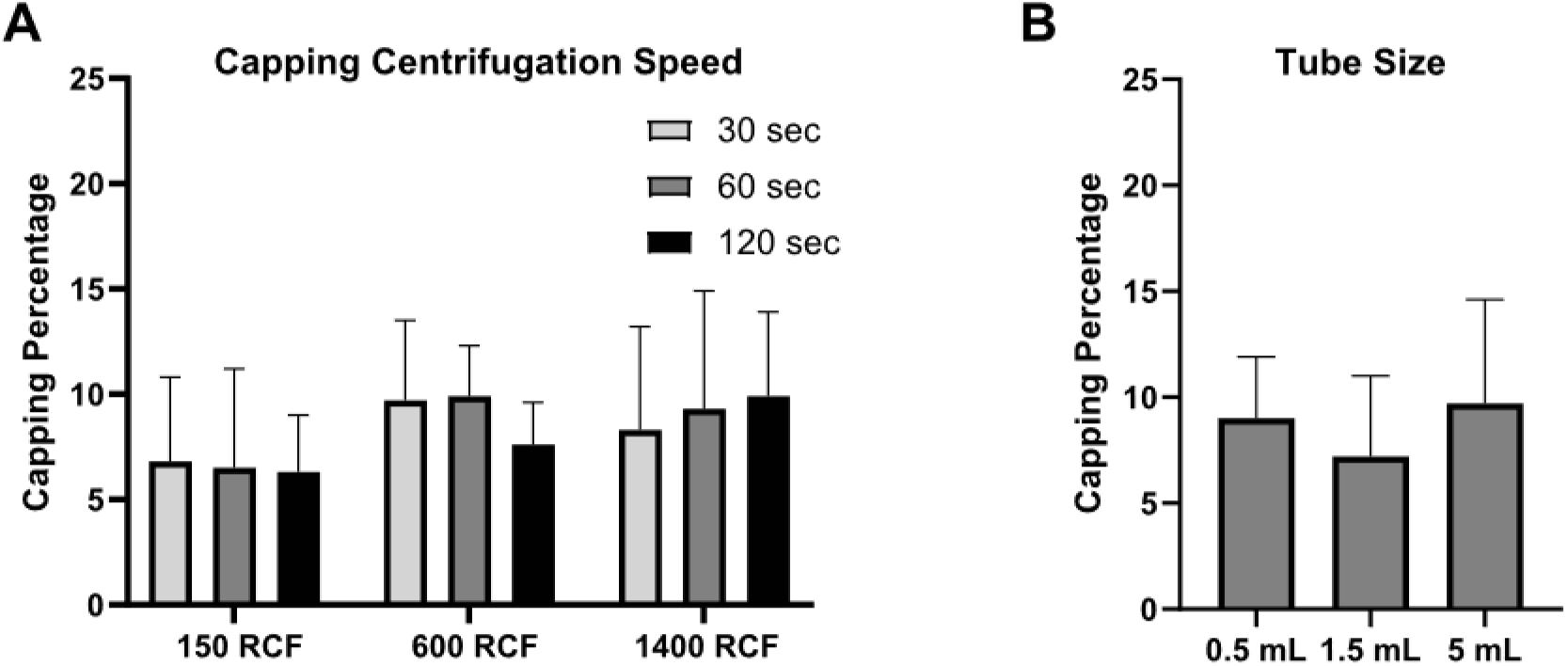
Capping centrifugation speed, tube size, and capping efficiency. Effect of **A)** centrifugation speed and duration and **B)** tube size on capped nanovial formation efficiency.

**Supplementary Figure 4.**
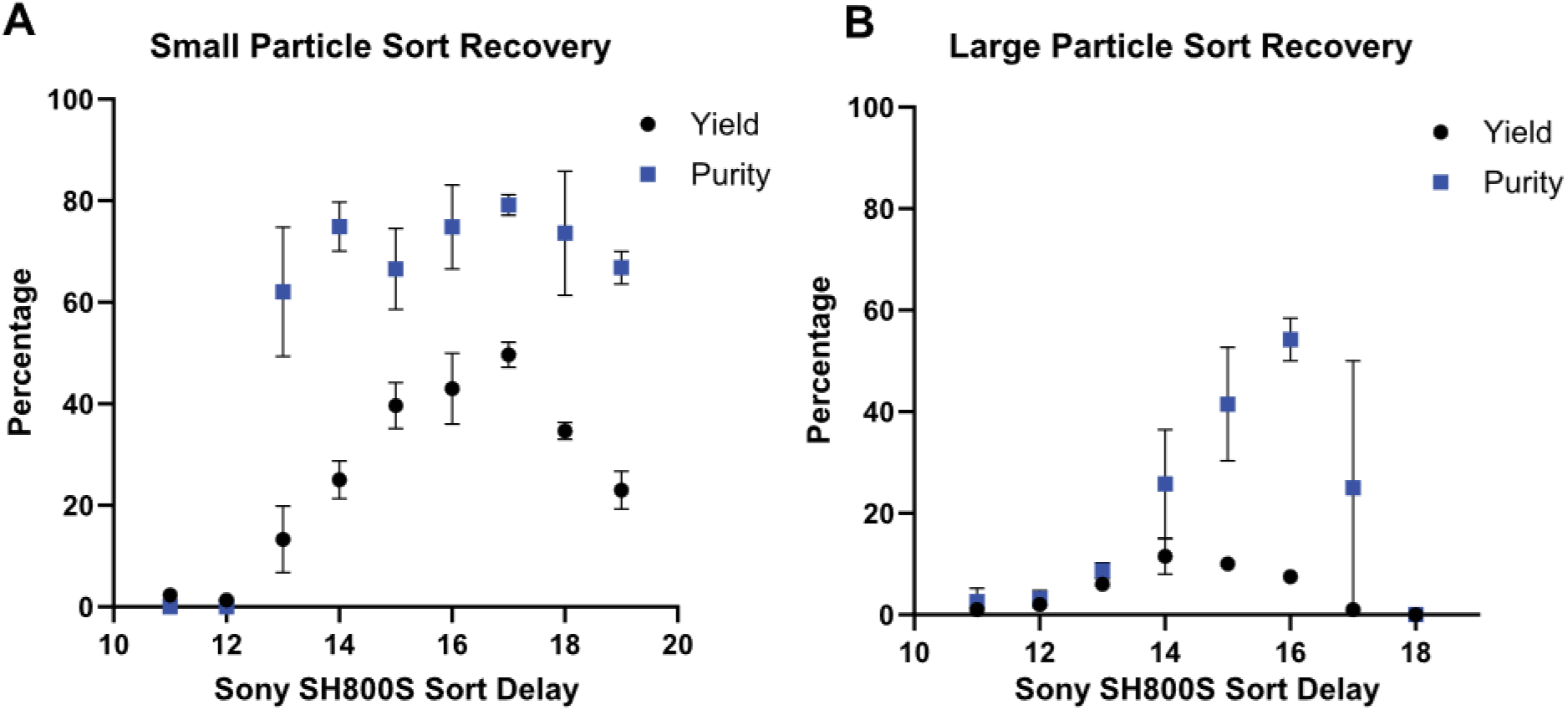
Flow cytometry sort yield and purity. **A)** Calculated capped nanovial sort yield and purity for different Sony SH800S sort delay settings using 40 µm nanovials with 30 µm capping particles. **B)** Capped nanovial sort yield and purity measurements for 70 µm nanovials with 40 µm capping particles. Yield is defined as fraction of capped nanovial sorted events that were visible by microscopy. Purity is defined as capped nanovial sorted events divided by all particles (capped nanovials, capping particles, and uncapped nanovials) imaged by microscopy.

**Supplementary Figure 5.**
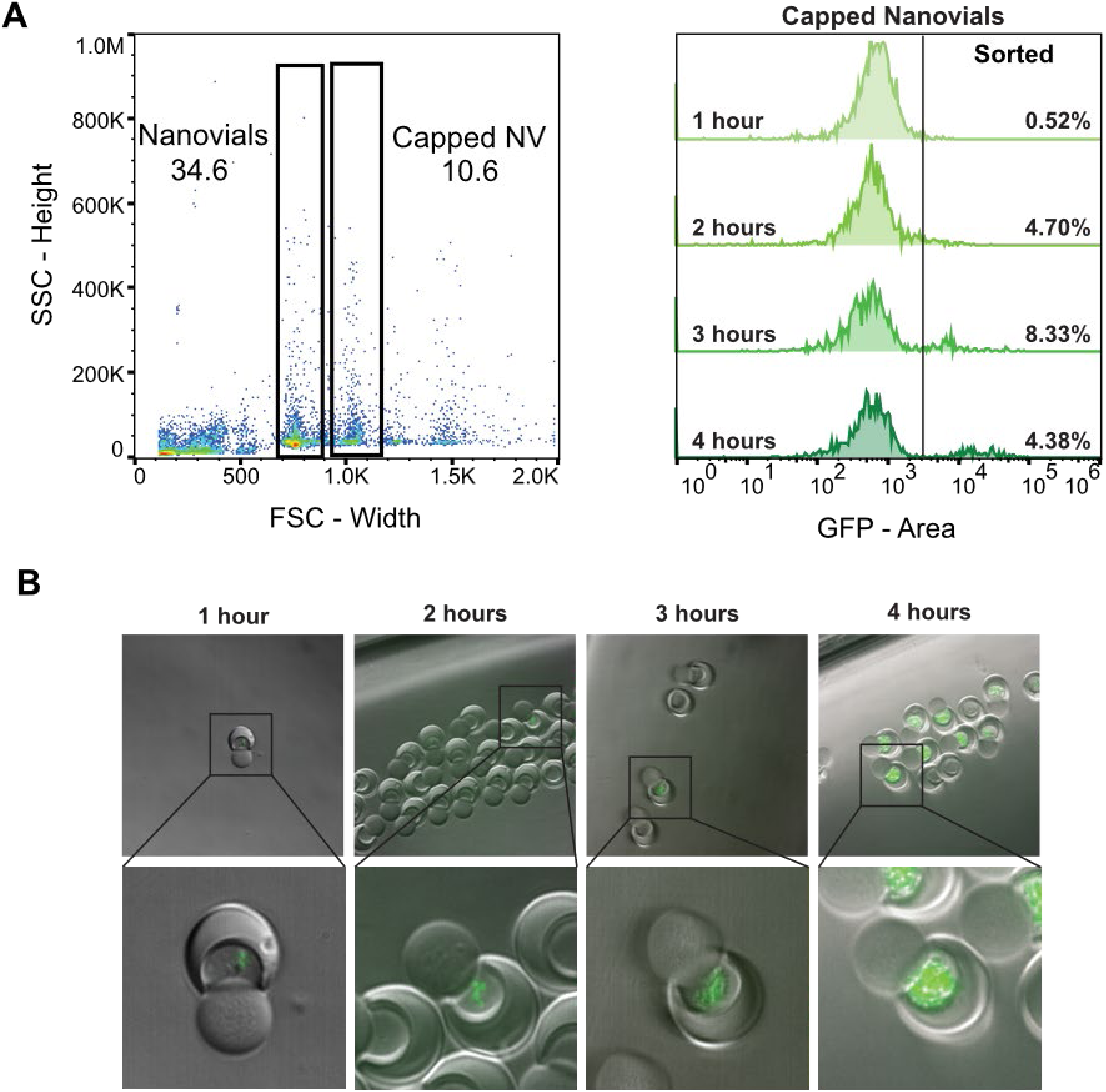
Enrichment of GFP-producing bacterial colonies. **A)** Gating strategy for enriching bacteria-loaded capped nanovials following different growth times. **B)** Fluorescence and brightfield microscopy of GFP-expressing *E. coli* following flow sorting following varying incubation times.

**Supplementary Figure 6.**
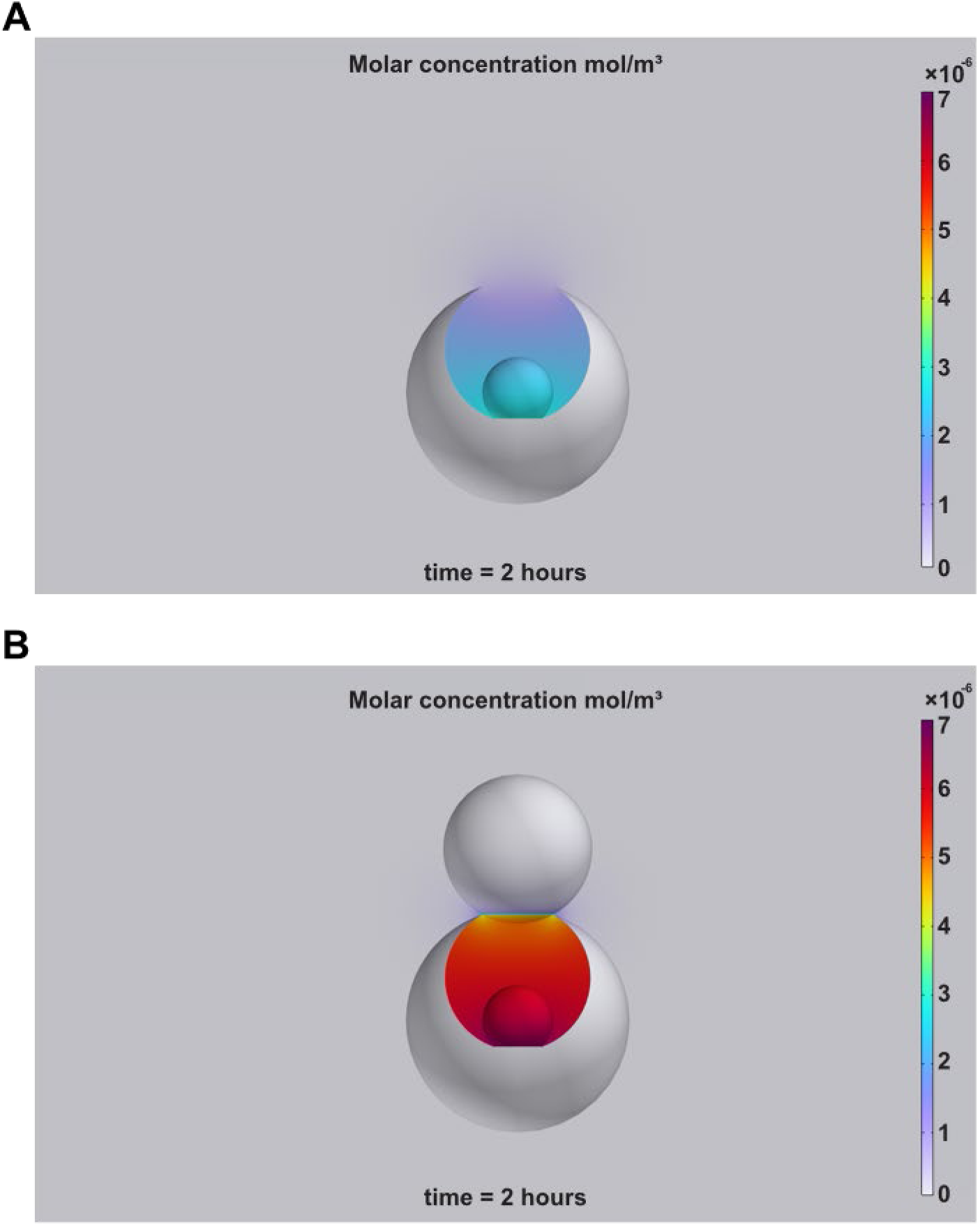
Simulation of antibody accumulation within nanovials and capped nanovials. COMSOL simulation results of secreted antibody accumulation in **A)** a nanovial and **B)** a capped nanovial.

**Supplementary Figure 7.**
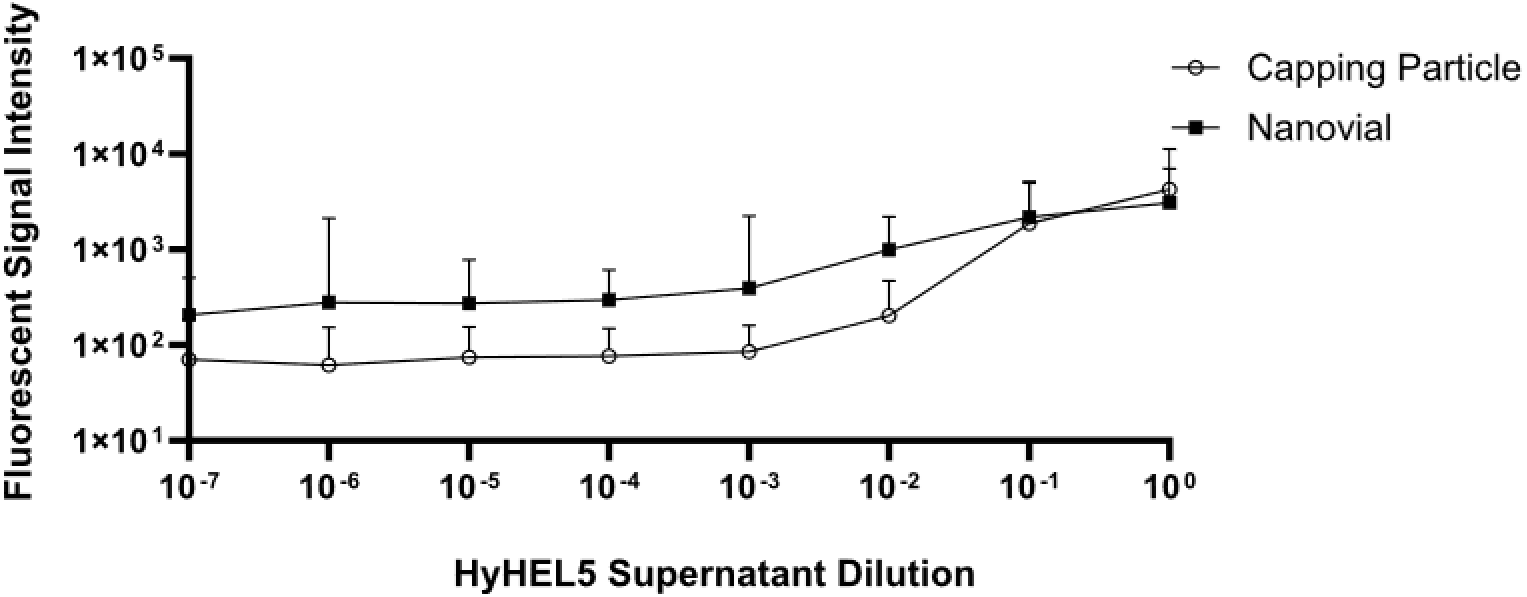
Particle dynamic range. Average capping particle and nanovial fluorescence intensity when incubated with varying dilutions of antibody-laden HyHEL5 supernatant.

**Supplementary Figure 8.**
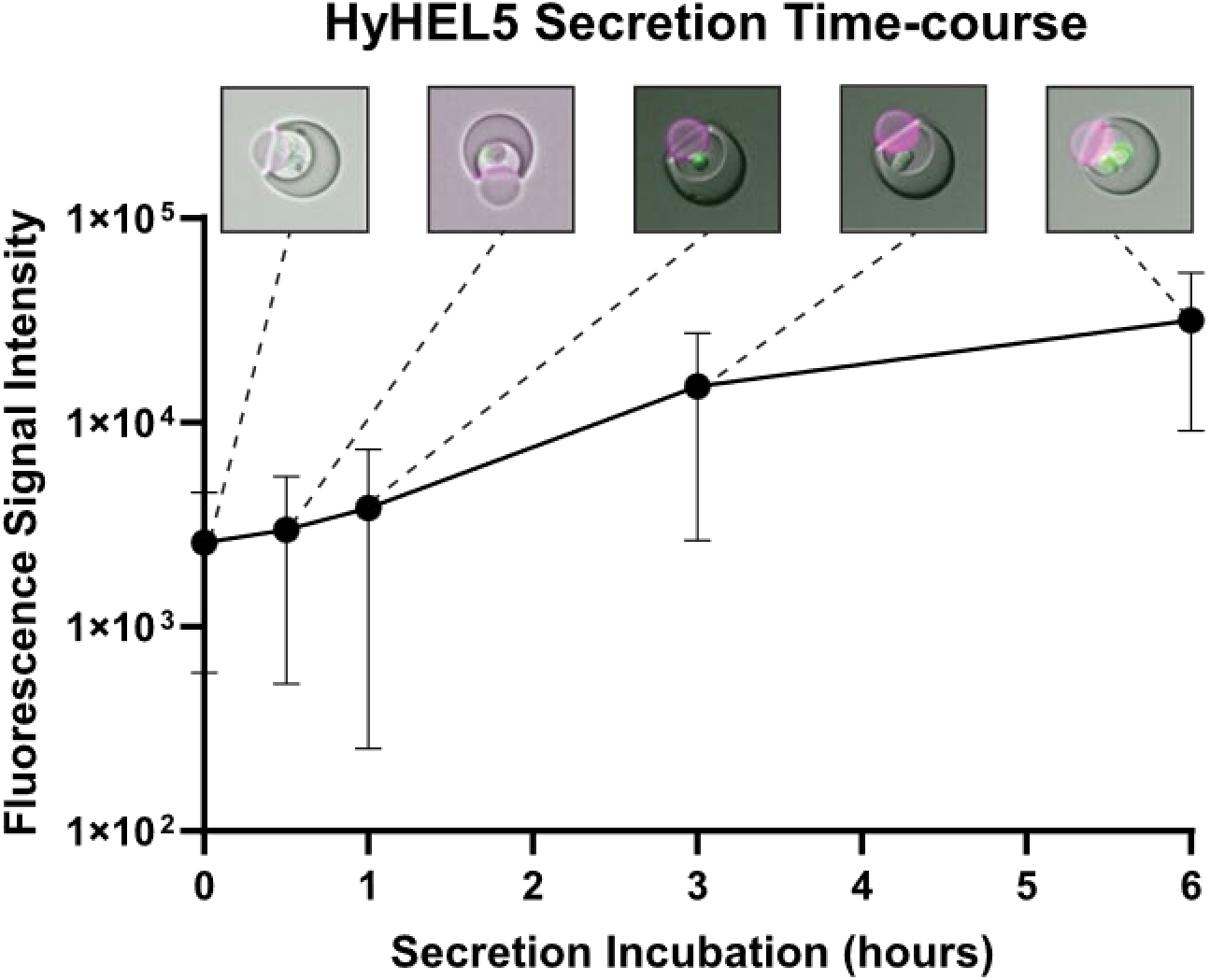
Single-cell secretion time-course. Average secretion signal on capped nanovials with increasing secretion incubation times. Brightfield/fluorescence overlaid images for selected time points.

**Video S1 Nanovial and capping particle settling dynamics.** Timelapse brightfield microscopy images of nanovials and capping particles settling in a well.

**Video S2 *S. cerevisiae* growth within capped nanovials.** Timelapse images of *S. cerevisiae* colony formation within capped nanovials over 16 hours.

**Video S3 *E. coli* growth within capped nanovials.** Timelapse images of *E. coli* expansion within capped nanovials over 5 hours.

