## Supplementary figures and images for "Sealable capped nanovials for high-throughput screening of cell growth and function"

### Video S3

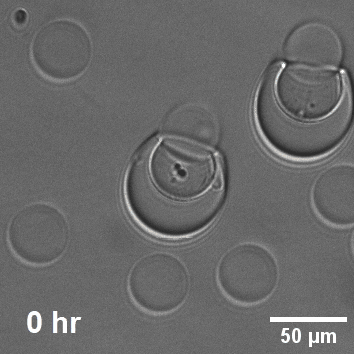
